# Molecular mechanism for strengthening E-cadherin adhesion using a monoclonal antibody

**DOI:** 10.1101/2022.03.06.483190

**Authors:** Bin Xie, Allison Maker, Andrew V. Priest, David M. Dranow, Jenny N. Phan, Thomas E. Edwards, Bart Staker, Peter J Myler, Barry M. Gumbiner, Sanjeevi Sivasankar

## Abstract

E-cadherin (Ecad) is an essential cell-cell adhesion protein with tumor suppression properties. The adhesive state of Ecad can be modified by the monoclonal antibody 19A11, which has potential applications in reducing cancer metastasis. Using x-ray crystallography, we determine the structure of 19A11 Fab bound to Ecad and show that the antibody binds to the first extracellular domain of Ecad near its primary adhesive motif - the strand-swap dimer interface. Molecular dynamics simulations and single molecule atomic force microscopy demonstrate that 19A11 interacts with Ecad in two distinct modes, one that strengthens the strand-swap dimer and one that does not alter adhesion. We show that adhesion is strengthened by the formation of a salt bridge between 19A11 and Ecad, which in turn stabilizes the swapped β-strand and its complimentary binding pocket. Our results identify mechanistic principles for engineering antibodies to enhance Ecad adhesion.

## Introduction

E-cadherin (Ecad) is an essential cell-cell adhesion protein that plays key roles in the formation of epithelial tissues and the maintenance of tissue integrity. Adhesion is mediated by the *trans* binding of Ecad ectodomains (extracellular regions) from opposing cell surfaces. Deficiencies in Ecad adhesion result in loss of contact inhibition and increased cell mobility ^1^, and are associated with the metastasis of gastric cancer ^2^, breast cancer ^3^, colorectal cancer ^4^ and lung cancer ^5^. Consequently, strategies that activate or strengthen Ecad adhesion may have potential applications in reducing cancer metastasis.

A powerful therapeutic approach that has been successfully used in regulating the binding of cell adhesion proteins are monoclonal antibodies (mAbs). For example, mAbs targeted against integrin adhesion proteins are used in the treatment of Crohn’s disease ^6-8^. Similarly, we have identified activating mAbs that target Ecad ectodomains and enhance cell-cell adhesion ^9^. In mouse models, one of these mAbs, 19A11, prevents the metastatic invasion of mouse lung cancer cells expressing human Ecad ^10,11^. In addition, we have shown that 19A11 can enhance Ecad’s epithelial barrier function and limit progression of inflammatory Bowel Disease (IBD) ^12^. Here, we resolve the molecular mechanisms by which mAb 19A11 strengthens Ecad adhesion.

We demonstrate that 19A11 strengthens adhesion by stabilizing strand-swap dimers, which are the predominant Ecad *trans* binding conformation. Strand-swap dimers are formed by the exchange of N-terminal β-strands (residues 1-12) between the outermost domains (EC1) of opposing Ecads. The exchange of β-strands results in the symmetric docking of a conserved anchor residue, tryptophan at the position 2 (W2), into a complementary pocket on the partner Ecad ^13,14^. Previous studies show that the two key structural and energetic determinants of Ecad strand-swap dimer formation are the stability of swapped β-strands ^15^ and their corresponding hydrophobic binding pockets ^16^. Using X-ray crystallography, molecular dynamics (MD) simulations, steered MD (SMD) simulations and single molecule atomic force microscopy (AFM), we show that 19A11 binding stabilizes both the β-strand and the hydrophobic pocket by forming key salt bridges. Our results identify the mechanistic principles underlying the activation of cadherin adhesion by mAbs.

## Results

### Crystal structure of 19A11 bound to Ecad

We co-crystallized the EC1-2 domains of human Ecad and 19A11 antibody fragment (Fab) and determined the structure at 2.2Å resolution (Fig. 1a, PDB ID code 6CXY). The structure refinement parameters are summarized in Supplementary Table 1. The crystal structure reveals that 19A11 recognizes two regions on the Ecad EC1 domain: residues 13-20, and residues 61-70. Binding of 19A11 does not cause gross conformational changes on Ecad and the root mean square deviation (RMSD) between crystal structures in the presence and absence (PDB ID code 2O72) of 19A11 is only 0.3Å. A closer look at the Ecad binding interface shows that binding of the antibody heavy chain and Ecad is primarily mediated by a salt bridge between K14 on Ecad and D58 on the 19A11 (Fig. 1b), and three Ecad:mAb hydrogen bonds (V48:R31, T63:Y104, and D64:G54). In addition, the crystal structure shows five hydrogen bonds between Ecad and the 19A11 light chain (K14:T100, F17:Y38, P18:N31, N20:S33, and K61:S33; Fig. 1c). The strong interaction between 19A11 and residues 13-20 of Ecad located at the base of the N-terminal β-strand led us to hypothesize that these interactions may strengthen Ecad strand-swap dimers.

**Fig. 1:**
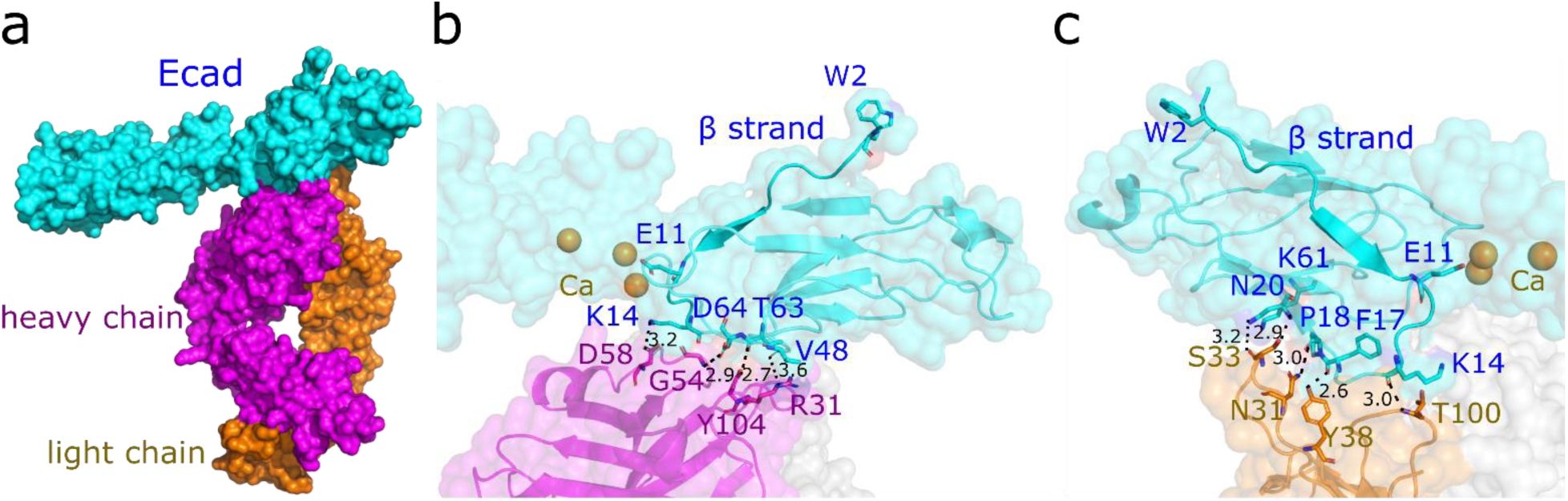
Structure of 19A11 bound to Ecad. (a) X-ray crystal structure of 19A11 Fab heavy chain (magenta) and light chain (orange) bound to Ecad EC1-EC2 domains (cyan). (b) Detailed view of the hydrogen bonds and salt bridges between 19A11 Fab heavy chain (magenta) and Ecad EC1 domain (cyan). (c) Detailed view of the interactions between 19A11 Fab light chain (orange) and Ecad EC1 domain (cyan). The distance between interacting atoms in (b) and (c) are shown in Å (black dashed lines).

### Molecular mechanisms of 19A11 mediated stabilization of strand-swap dimers

To resolve the detailed molecular interactions between 19A11 and Ecad, we performed MD simulations on three different structures: (*i*) Ecad strand-swap dimer *(EcadA and EcadB)* without 19A11 Fab (**0ab**, Fig. 2a, green structure, PDB ID code 2O72), (*ii*) Ecad strand-swap dimer bound to a single 19A11 Fab *(abA bound to EcadA)* (**1ab**, Fig. 2b, PDB ID code 6CXY), and (*iii*) Ecad strand-swap dimer bound to two 19A11 Fabs *(abA bound to EcadA and abB bound to EcadB)* (**2ab**, Fig. 2c). In the 0ab, 1ab, and 2ab conditions, we performed five independent simulations. Every MD simulation was performed for 60ns, which was long enough for the RMSD relative to the structures at the start of the simulations to stabilize (Supplementary Fig. 1).

**Fig. 2.**
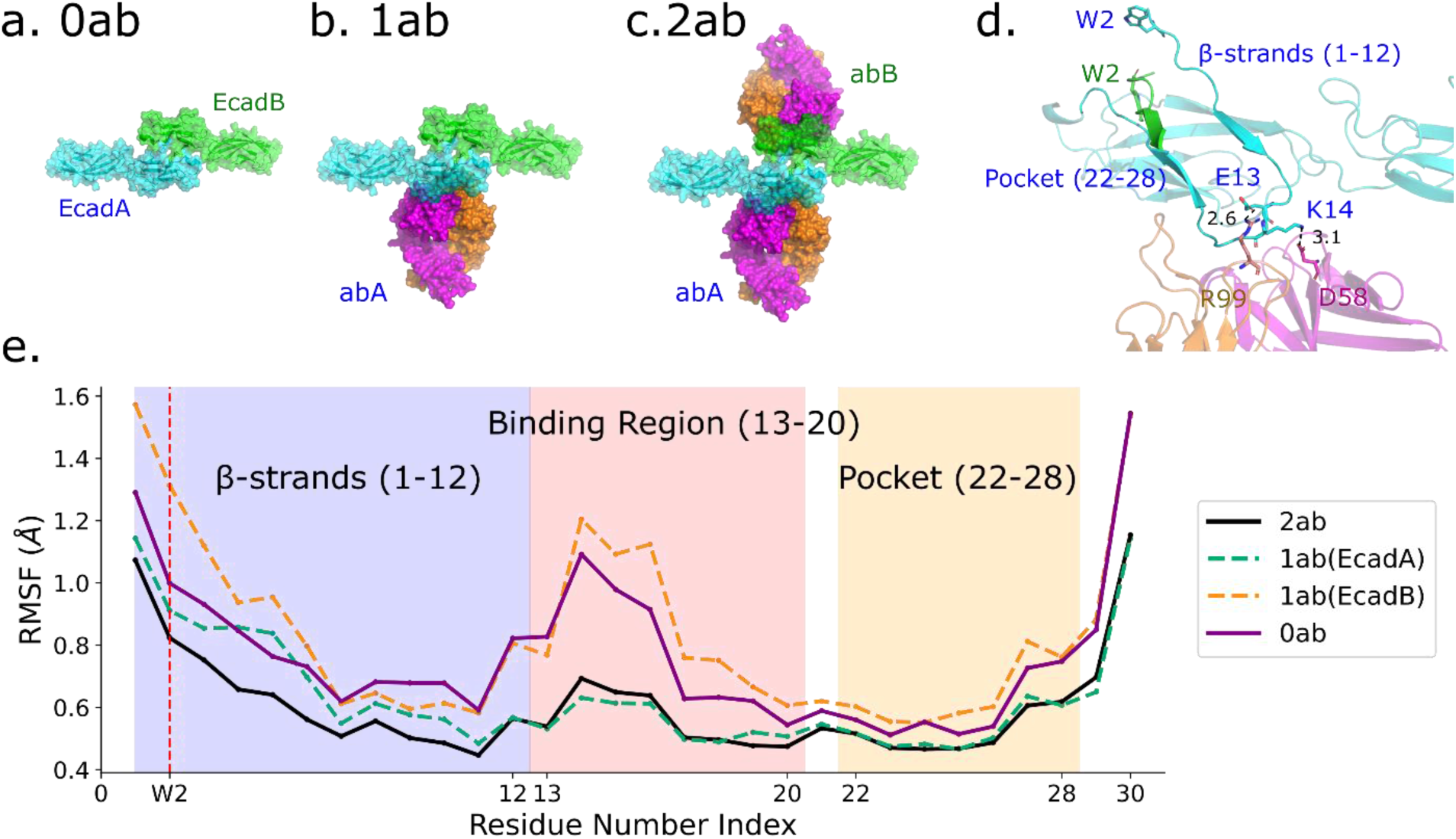
Binding of 19A11 stabilizes both the Ecad β-strand and the W2 hydrophobic pocket. MD simulations were performed with (a) Ecad strand-swap dimer (EcadA and EcadB) in the absence of 19A11 (0ab), (b) Ecad strand-swap dimer with a single 19A11 Fab (abA) bound to EcadA (1ab), and (c) Ecad strand-swap dimer with two 19A11 Fabs (abA and abB) bound to both Ecads (2ab). (d) Ecad-antibody binding interface. Two salt bridges are observed: E13-R99 and K14-D58. The 19A11 binding region is located between the β-strands and the W2 hydrophobic pockets (referred to as ‘pocket’) on Ecad. (e) Average RMSF values for residues 1-30 of Ecad in the 2ab case (solid black), EcadA in the 1ab case (dashed green), EcadB in the 1ab case (dashed orange), and Ecad in the 0ab case (solid purple). The W2 position is highlighted using a vertical dashed red line. The lower RMSF values show that binding of 19A11, stabilizes the β-strand and the W2 hydrophobic pocket of both Ecads in the 2ab case, while it only stabilizes EcadA in the 1ab case, which is bound to 19A11.

Since the β-strand and its complementary binding pocket are essential components of Ecad strand-swap dimers (Fig. 2d), we first tested if there was a change in the stability of either region upon 19A11 binding. We tested the stability by measuring the root-mean-square fluctuations (RMSF, the standard deviation of the atomic positions) of the corresponding α-carbon residues during the last 10 ns of all MD simulations (Fig. 2f). The average RMSFs for EcadA and EcadB in the 1ab conditions were calculated separately since only EcadA is bound to an antibody (Fig. 2f, dashed green and dashed orange lines). The average RMSF for Ecads in the presence and absence of the antibody showed that 19A11 binding reduces the RMSF of its corresponding binding regions on Ecad (residues 13-20). Binding of 19A11 also stabilizes the two adjoining regions, the β-strand (residues 1-12) and the partial complementary pocket (residues 22-28), which are essential for the strand-swap dimer formation.

Next, we examined specific interactions between 19A11 and Ecad to determine the molecular mechanisms by which mAb binding stabilizes the β-strand and pocket region. We focused on two salt bridges that form between 19A11 and Ecad, downstream of the β-strand (Fig. 2d). The first salt-bridge occurs between E13 on Ecad and R99 on the 19A11 light chain while the second salt-bridge forms between K14 on Ecad and D58 on the 19A11 heavy chain (Fig. 2d). We measured the distances between the salt bridge charges in every MD simulation, in both the 2ab (Fig. 3a-e) and 1ab (Supplementary Fig. 2) conditions, from 20 ns to 60 ns. Based on the criterion that salt bridges form when the median distance between charged atoms is less than 4Å, we concluded that in 40% of the 2ab simulations (sets 1 and 2, Fig. 3a-b), both Ecads form at least one salt bridge with the bound 19A11. However, in the remaining 60% of the simulations, only one of the Ecad formed a salt bridge with 19A11 (EcadB of set 3, EcadB of set 4, and EcadA of set 5; Fig. 3c-e). Importantly, the salt bridges formed by sets 3-5 in the 2ab simulations were similar to the 1ab condition (Supplementary Fig. 3), in that only one Ecad formed a salt bridge (since there was only one 19A11 Fab present in the 1ab condition).

**Fig. 3:**
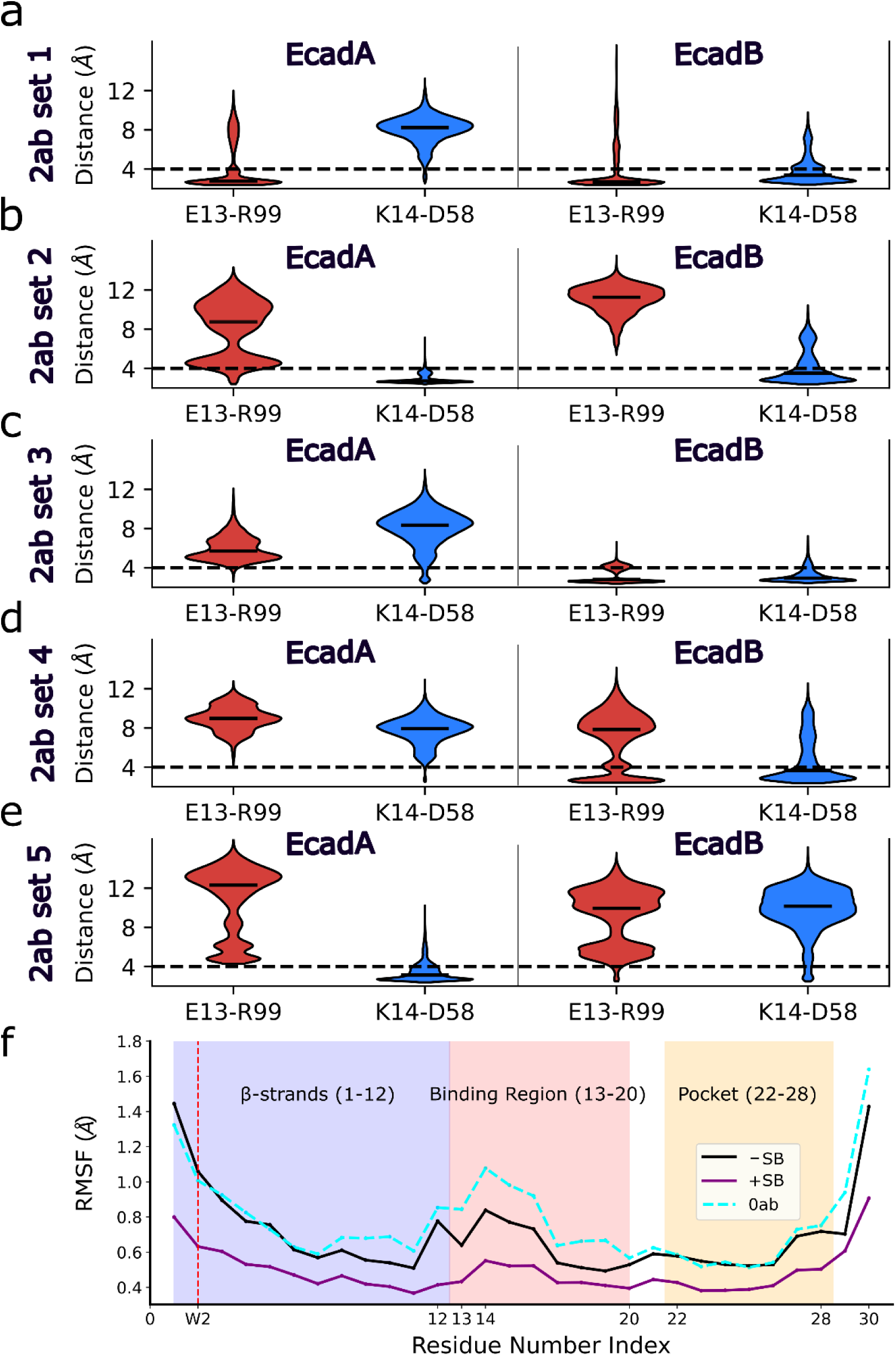
Salt bridges between 19A11 and Ecad stabilize the β-strand and the W2 hydrophobic pocket. (a-e) Violin plots of the distances between charged atoms in the E13-R99 and K14-D58 salt bridges measured during the last 40 ns of each 2ab MD simulation. The median distance is shown as a black line on each violin. Distance for EcadA and EcadB are shown in the left and right panels respectively. Distances measured for E13-R99 interactions during the MD simulations are shown in red while charged atoms distances for K14-D58 are shown in blue. (a) simulation-1 (set 1); (b) simulation-2 (set 2); (c) simulation-3 (set 3); (d) simulation-4 (set 4); (e) simulation-5 (set 5). Both EcadA and EcadB form at least one salt bridge with the bound 19A11 in set 1 and set 2. However, only one of the Ecad formed a salt bridge with 19A11 in sets 3-5 (EcadB in set 3, EcadB in set 4, and EcadA in set 5). (f) Comparison of the average backbone RMSF values when both interacting Ecads form at least one salt bridge with 19A11 (purple solid line), when both Ecads do not form a salt bridge (black solid line), and in the absence of 19A11, *i*.*e*. 0ab (dashed cyan line). The RMSF for W2 is highlighted using a vertical dashed red line. The β-strand and the W2 hydrophobic pocket have a lower RMSF when salt bridges are formed as compared to when no salt bridges are formed.

When all Ecads form at least one salt bridge with their corresponding 19A11, the average RMSF of the β-strand and pocket region was lower (Fig. 3f, purple line), demonstrating that salt bridge formation stabilizes both the β-strand and pocket region. In contrast, the stability of the β-strand and pocket region when all Ecads do not form a salt bridge (Fig. 3f, black line), was approximately the same as the 0ab condition (Fig. 3f, dashed cyan line). Based on these results we concluded that 19A11 can interact with Ecad in two different modes: one which stabilizes the β-strand and the pocket region by forming either E13-R99 or K14-D58 interactions, and a second mode which does not stabilize the β-strand and the pocket region because the salt bridges are not formed.

### 19A11 mediated stabilization of strand-swap dimer leads to stronger Ecad adhesion

Since each pocket region forms hydrogen bonds with a β-strand on its partner Ecad ^16^, we hypothesized that stabilizing the pocket and β-strand strengthens adhesion by retaining each β-strand in its swapped position and keeping each W2 inserted into its opposing pocket. To test this hypothesis, we performed SMD simulations. We fixed the C-terminal end of one Ecad in the final structure of each MD simulation and pulled on the other Ecad C-terminus with a constant force of ∼665 pN (supplementary movie 1). During each SMD simulation, we measured the interfacial binding area between the two Ecads, which we estimated using the change in the solvent accessible surface area (ΔSASA) ^17^; decrease of ΔSASA to zero corresponded to the rupture of the interacting *trans* dimer. In the 0ab condition (Fig. 4a), the ΔSASA dropped to zero at ∼900 ps while in the 1ab condition (Fig. 4b) the interactions between Ecads lasted marginally longer and broke at ∼1000 ps. In contrast to the 0ab and 1ab conditions, we measured two populations in the 2ab condition: one that remained bound for a longer time, and another that unbound on a time scale closer to the 0ab and 1ab conditions. Ecads in sets 1 and 2 of the 2ab condition interacted for ∼2700 ps suggesting a strong bound state, while in sets 3-5 the interactions only lasted for ∼1600 ps (Fig. 4c).

**Fig. 4:**
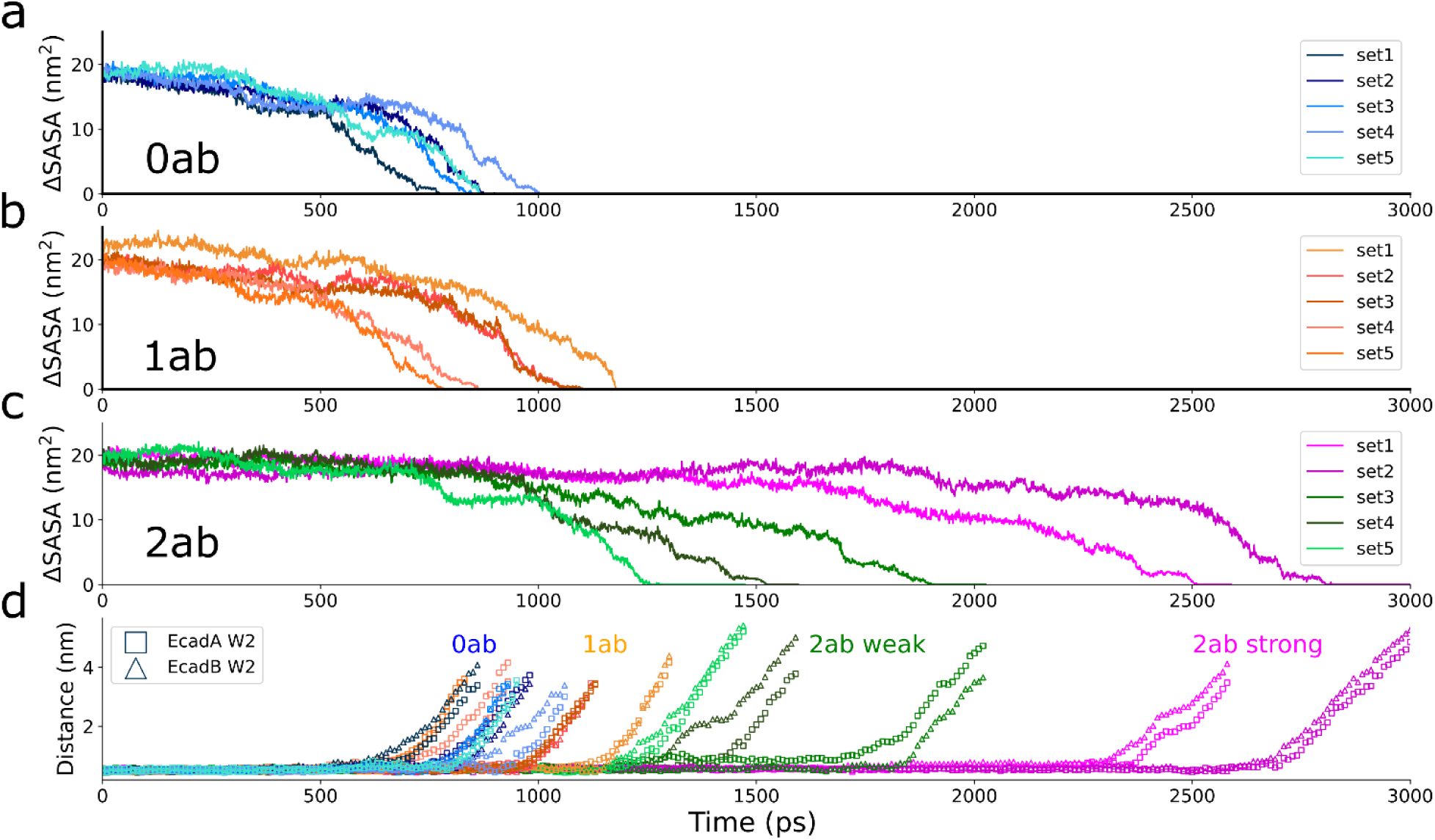
Adhesion strengthening requires two bound 19A11 antibodies to form salt bridges with partner Ecads. Constant force SMD simulations with change in Ecad-Ecad interfacial area calculated from the change in the solvent accessible surface area (ΔSASA), in the (a) 0ab condition, (b) the 1ab conditions, and (c) the 2ab conditions. (d) Distance between center of mass of W2 and the center of mass of the hydrophobic pockets in each of the constant force SMD simulations. While the lifetimes of the Ecad-Ecad bonds are similar in the 0ab, 1ab and sets 3-5 of the 2ab condition, the lifetime of the Ecad-Ecad bond in the sets 1-2 of the 2ab condition, where both interacting Ecads form at least one salt bridge with 19A11, are substantially longer *(also see supplementary video 1)*.

As an additional measure of strand-swap dimer stability, we calculated the distances between the center of mass of W2 and the center of mass of its complimentary binding pocket during the constant force SMD (Fig. 4d). These measurements tell us how long W2 is retained in the hydrophobic pocket because the β-strands remain in a swapped position. Similar to the ΔSASA measurement, W2 in the 0ab condition, 1ab condition, and sets 3-5 of the 2ab condition exited its complimentary pocket much earlier than 2ab set 1 and 2 (Fig. 4d).

Taken together, our simulations show that formation of salt bridges between two 19A11 Fabs and their corresponding Ecads in a strand-swap dimer (*i*.*e*. sets 1 and 2 of the 2ab condition), stabilize the β-strand and pocket and strengthen adhesion. In contrast, when a salt bridge is formed between 19A11 and only one Ecad in a strand-swap dimer, the β-strand and pocket region are less stable and adhesion is not strengthened.

### 19A11 strengthens Ecad interactions at the single molecule level

To experimentally validate our simulations, we directly tested the effect of 19A11 binding on the strengthening of single Ecad interactions using AFM force measurements. We immobilized the complete extracellular region of canine Ecad (EC1-5) on an AFM cantilever and glass substrate (Fig. 5a), and measured Ecad-Ecad interactions in the presence and absence of 19A11 Fab (Fig. 5a-c). We have previously demonstrated that 19A11 Fab strengthens adhesion of Madin-Darby canine kidney (MDCK) cells expressing canine Ecad ^9^. Furthermore, the human Ecad and canine Ecad amino acid sequences are conserved with 91% sequence identity in the EC1 domain (Supplementary Fig. 3). However, since we performed AFM experiments with recombinant canine Ecad ectodomains, we independently verified 19A11 Fab binding using western blotting (Supplementary Fig. 4).

**Fig. 5:**
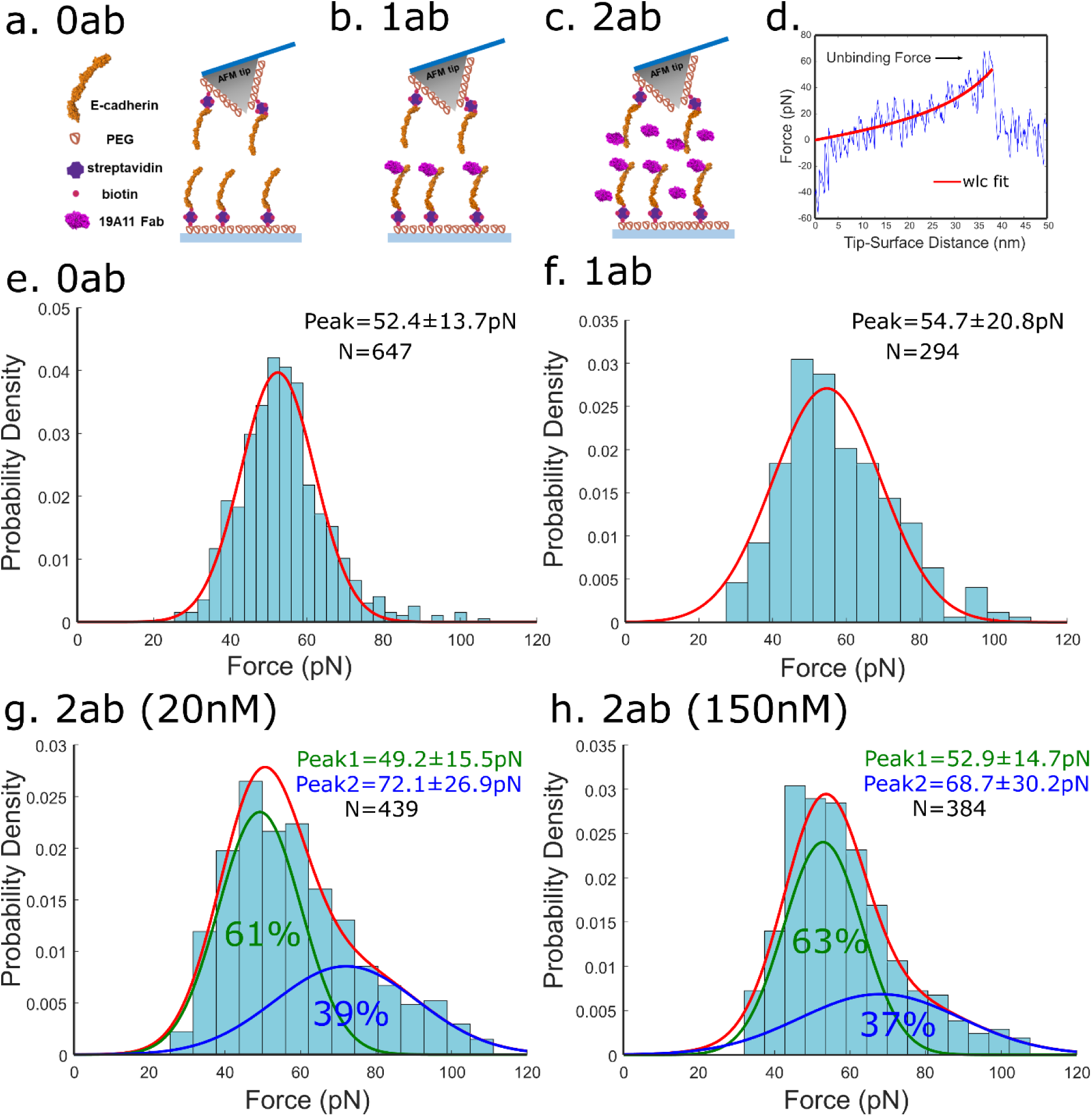
Direct, single molecule measurements of 19A11 mediated strengthening of Ecad homophilic adhesion. (a) Scheme for AFM experiment carried out in the absence of 19A11 (0ab). Ecads were immobilized on an AFM tip and substrate functionalized with polyethylene glycol (PEG) tethers. (b) Scheme for AFM force measurements with one antibody (1ab). Only the substrate was incubated with 19A11. (c) Scheme for two antibody AFM experiment (2ab). Both the AFM tip and substrate were incubated with 19A11 (2ab). (d) Example force curve. Stretching of the PEG tether, which served as a ‘signature’ of a single molecule unbinding event, was fit to a worm-like chain model (red line). (e) Probability density of Ecad-Ecad unbinding forces measured in the absence of 19A11. Forces are Gaussian distributed (red line) with a peak force of 52.4±13.7 pN. (f) Probability density of unbinding forces measured in the 1ab condition fitted to a Gaussian distribution (red line) with peak=54.7±20.8 pN. (g) Probability density of Ecad-Ecad unbinding forces in the presence of 20 nM 19A11 fit to a bimodal Gaussian distribution. While the first peak at 49.2±15.5 pN (green line) corresponds to the ‘native’ Ecad unbinding force, the second peak at 72.1±26.9 pN (blue line) corresponds to strengthened adhesion. (h) Increasing the concentration of 19A11 in solution to 150 nM, yields a similar bimodal Gaussian distribution with peaks at 52.9±14.7 pN (green line), and 68.2±30.2 pN (blue line). This demonstrates that the bimodal distribution of forces does not occur due to low 19A11-Ecad binding affinity but rather because 19A11 binds to Ecad in two distinct modes.

A typical AFM measurement consisted of bringing the cantilever and substrate—both functionalized with Ecad—into contact and allowing the opposing cadherins to interact. The tip was then withdrawn from the substrate at a constant velocity, and the force required to rupture the Ecad-Ecad bond was measured. Interaction of opposing Ecads resulted in unbinding events that were characterized by the non-linear stretching of PEG tethers, which were fit to a worm-like chain model (WLC) using least squares fitting (Fig. 5d).

Analogous to the MD simulations, AFM experiments were performed under three conditions: “0ab”, “1ab”, and “2ab”. AFM measurements without 19A11 Fab (0ab; Fig. 5a) were performed to verify Ecad-Ecad binding and served as a control to benchmark 19A11 strengthening. For the 1ab experiment, we incubated only Ecad immobilized on the glass coverslip with 19A11 Fab (Fig. 5b). Incubating both the coverslip and the cantilever with 19A11 and performing the AFM experiments in the presence of free 19A11 Fab in the measurement buffer, constituted the 2ab condition (Fig. 5c).

To quantitatively compare binding strengths, we performed all AFM measurements at a single pulling velocity (1μm/s). Unbinding forces between Ecad-Ecad bonds without 19A11 bound (0ab) showed a single Gaussian distribution and yielded an unbinding force of 52.4 ± 13.7 pN (Fig.5e). As predicted by the MD simulations, no strengthening was also observed for 1ab experiments (peak force = 54.6 ± 20.8 pN; Fig. 5f). However, when the experiments were performed in the presence of 20 nM 19A11, which corresponds to mAb bound to both Ecads (2ab), the unbinding force distribution was bimodal with one peak corresponding to strengthened Ecad-Ecad bonds (72.1 ± 26.9 pN; Fig. 5g) and one that was comparable to the 0ab and 1ab conditions (49.2 ± 15.5 pN; Fig. 5h). As predicted by the MD and SMD simulations, only ∼40% of Ecad interactions were strengthened (blue peak; Fig. 5g).

To confirm that the bimodal distribution of unbinding forces was due to 19A11 binding in two distinct modes (as predicted by the simulations), and not due to low affinity of Ecad for the mAb, we increased the concentration of the 19A11 Fab to 150 nM in the 2ab experiments. Again, we measured Ecad strengthening, but the unbinding forces still showed only ∼40% of Ecads that were strengthened (peak force = 68.7 ± 30.2 pN; Fig. 5h). Thus, in excellent agreement with the simulations, our AFM results showed that while the binding of a single 19A11 Fab to Ecad did not enhance adhesion, the binding of two 19A11 Fabs, strengthened ∼40% of Ecad interactions.

## Discussion

It has previously been shown that a key factor for strand-swap dimer formation is the stabilization of the swapped β-strand (residues 1-12) ^15^. Previous studies also show that residues 22-28 within the hydrophobic pocket enhance Ecad *trans* dimerization affinity by forming hydrogen bonds, specifically Asp1-Asn27 and Val3-Lys25, with the β-strand of its partner Ecad ^16^. Our X-ray crystal structure demonstrates that 19A11 mAb binds to the EC1 domain, between the β-strand and the pocket region of Ecad. Using MD simulations and AFM force measurements, we show that 19A11 forms two key salt bridge interactions with Ecad which stabilizes both the swapped β-strand and the pocket region which houses a Trp2 from its binding partner. Consequently, to strengthen Ecad adhesion, one of these salt bridges needs to be formed between both Ecads in in the *trans* dimer and their bound 19A11. Due to the stochastic formation of salt bridges, 19A11 can interacts with Ecad in two distinct modes: one that strengthens the Ecad-Ecad bond and one that does not change its adhesion.

In addition to forming robust strand-swap dimers, Ecads also adhere in a weaker X-dimer conformation, which is mediated by a salt-bridge between K14 and D138 on the opposing Ecads ^18,19^. Ecad ectodomains are believed to initially form X-dimers and then transition to a strand-swap dimer conformation ^19-21^. While our crystal structure shows that K14 on Ecad forms a salt bridge with D58 on the 19A11 heavy chain, it also demonstrates that the Ecads interact in a strand-swap dimer conformation. It is possible that the salt bridge between Ecad and 19A11 forms after strand-swap dimerization has occurred. Alternately, it is possible that K14 can form salt bridges with both D58 (on 19A11) and D138 (in an X-dimer) simultaneously (or in resonance) and initiate strand-swap dimer formation. Finally, it is also possible that strand-swap dimer formation occurs via alternate pathways that are independent of X-dimers, as has been observed in cell-free biophysical measurements ^19,21,22^. Irrespective, our measurements do not probe how the strand-swap dimers form, but instead demonstrate that antibody-bound strand-swap dimers are strengthened upon formation of key salt bridge interactions.

Besides constant force SMD simulations, we mimicked the single molecule AFM experiments *in silico* by performing SMD simulations at a constant pulling velocity (0.005 nm/ps, Supplementary Fig. 5). While the unbinding forces in the AFM experiments and SMD simulations cannot be directly compared because of their different pulling velocities, the maximum forces measured in the simulations served as a qualitative benchmark of the strength of the Ecad *trans* dimer. As anticipated from the AFM experiments, our constant-velocity SMD simulation data showed that the average maximum forces observed in the 0ab, 1ab, and weaker 2ab (sets 3-5) conditions were comparable. In contrast, the stronger 2ab (sets 1, 2) conditions had a higher average maximum force.

We have previously shown that 19A11 binding to Ecad ectodomains in Colo 205 cells induces p120-catenin dephosphorylation which correlates with adhesion activation ^9^. While the current study does not investigate these intracellular consequences of 19A11 binding, we show that 19A11 also strengthens Ecad ectodomain adhesion, independent of the cytoplasmic region.

Our work establishes novel principles for design of mAbs to enhance cadherin adhesion. Unlike integrin activating antibodies which regulate ectodomain conformation ^6^, we show that 19A11 strengthens adhesion, not by inducing gross conformational changes in the Ecad ectodomain, but rather by selectively stabilizing the swapped β-strand and its complimentary binding pocket. Consequently, our results demonstrate that selectively targeting these structural and energetic determinants of strand-swap dimer formation are sufficient to strengthen Ecad adhesion. We anticipate that this approach may prove to be a useful strategy in designing activating mAbs for other classical cadherins which interact by N-terminal β-strand swapping.

## Methods

### Purification of human Ecad EC1-2 for X-ray crystallography

We used residues 155-371 to encompass EC1-2 of the human Ecad extracellular domains; the signal sequence and pro-domain were deleted (Δ1-154). EC1-2 was expressed as a fusion protein by attaching 6x His tagged SMT3 to the N-terminus ^23^. The EC1-2 construct was cloned into pET21a plasmid system and transformed into BL21 (DE3) competent cells (Novagen). Cultures were grown in autoinduction media ^24^ overnight and harvested via centrifugation. Thawed bacterial pellets were lysed by sonication in 200 ml buffer containing 25 mM HEPES pH 7.0, 500 mM NaCl, 5% Glycerol, 0.5% CHAPS, 10mM Imidazole, 10 mM MgCl_2_, and 3 mM CaCl_2_. After sonication, the crude lysate was clarified with 2μl (250 units/μl) of Benzonase and incubated while mixing at room temperature for 45 minutes. The lysate was then clarified by centrifugation at 10,000 rev min^−1^ for 1 h using a Sorvall centrifuge (Thermo Scientific) followed by filtration via 0.45μm syringe filters. The clarified supernatant was then passed over a Ni-NTA His-Trap FF 5 ml column (GE Healthcare) which was pre-equilibrated with loading buffer composed of 25 mM HEPES pH 7.0, 500 mM NaCl, 5% Glycerol, 20 mM Imidazole, and 3 mM CaCl_2_. The column was washed with 20 column volumes (CV) of loading buffer and was eluted with loading buffer plus 500 mM imidazole in a linear gradient over 10 CV. Peak fractions, as determined by UV at 280 nm, were pooled and concentrated to 10mL. Pooled fractions were dialyzed overnight against 4 liters buffer containing 500mM NaCl, 25mM HEPES, 5% Glycerol, 3mM CaCl_2_ (SEC Buffer) with His-tagged Ulp1 protease added to cleave the 6xHis-SMT3 fusion protein at a ratio of 1 mg Ulp1 for 1000 mg protein. Dialysate was passed over a Ni-NTA His-Trap FF 5 ml column to remove 6xHis-SMT3 fusion protein and Ulp1. Flow-through from the nickel column was concentrated to 5 ml and passed over a Superdex 75 SEC column (GE) equilibrated with SEC Buffer. The peak fractions were collected and analyzed for the presence of the protein of interest using SDS–PAGE. The peak fractions were pooled and concentrated using Macrosep 20 mL 10K MWCO protein concentrators (Pall). Aliquots were flash-frozen in liquid nitrogen and stored at −80°C until use for preparation of hE-cadherin EC1-2-Activating Fab complexes.

### Purification of 19A11 Fab and Fab-Ecad complexes for X-ray crystallography

Sequences coding for the heavy chain of the 19A11 Fab fragment were cloned into pcDNA3.4 with a Twin-Strep tag added after the C-terminal residue (SAWSHPQFEKGGGSGGGSGGGAWSHPQFEK*). ExpiCHO cells (ThermoFisher) were transfected with the appropriate light chain and heavy chain encoding plasmids for each Fab following the ExpiFectamine CHO Transfection Kit (ThermoFisher) high titer protocol. To obtain a single pure species of 19A11 for crystallography, a minor glycosylated product (10% of the total) was removed by incubating with ConA slurry (GE Healthcare) for 4 hours at 4°C on a rotator. Purification of StrepTag Fabs from ExpiCHO culture medium was performed using StrepTactin XT Superflow High Capacity resin (IBA), elution with 50 mM biotin, followed by buffer exchange with PD-10 columns to 50 mM Tris pH 8.0, 0.15 M NaCl and 3 mM CaCl_2_. Isolation of a single species for each Fab was verified by PAGE and activation of cellular Ecad was confirmed by Colo205 activation assay ^9^. hEC1-2 was incubated with a 1.6x molar excess of conA-purified 19A11-Fab and incubated overnight at 4°C. Complex was purified with SEC with a Superose 6 10/300 GL column and concentrated to 10.3 mg/ml in 50 mM Tris, 150 mM NaCl, 3 mM CaCl_2_, pH 8.0.

### Crystallization of human Ecad with 19A11 Fab

Protein complex (0.1 μL) was mixed with 0.1 μL of crystallization solution (Wizard 3/4, well H12, Rigagku Reagents) containing 15% (w/v) PEG-20000, 0.1 M HEPES/NaOH (pH 7.0), and equilibrated against 50 μL of crystallization solution in an MRC2 vapor diffusion tray (SWISSCI). Crystals were harvested and cryoprotected with crystallization solution supplemented with 20% ethylene glycol and flash frozen in liquid nitrogen.

### Data collection and structure solution

Ecad with 19A11 diffraction data were collected at the Life Sciences Collaborative Access Team beamline 21-ID-F at the Advanced Photon Source (Argonne National Laboratory) on a Rayonix MX300 CCD detector at a wavelength of 0.97872 Å. Data were indexed and integrated with XDS and scaled with XSCALE ^25^. The structure was solved with Phaser ^26^ using PDB 2O72 as a search model for Ecad and PDB 4WEB as a search model for 19A11. The model was refined with iterative rounds of refinement with Phenix ^27^ and manual model building in Coot ^28,29^. The quality of the structure was checked with Molprobity ^30^.

### Purification of canine Ecad ectodomains for AFM

Generation of Ecad monomer plasmids containing a C-terminal Avi tag has been described previously ^31^. The plasmids were incorporated into pcDNA3.1(+) vectors and were transiently transfected into HEK 293T cells using PEI (Milipore Sigma) as previously described ^32^. Three days post transfection, conditioned media was collected for protein purification. Purification of Ecad were performed using methods described previously ^33-35^. Media containing his-tagged Ecads was passed through a chromatography column containing Ni-NTA agarose beads (Qiagen). Beads were then washed with a pH 7.5 biotinylation buffer (25 mM HEPES, 5 mM NaCl, and 1 mM CaCl_2_). Ecads bound to the Ni-NTA beads were biotinylated with BirA enzyme (BirA 500 kit; Avidity) for 1 h at 30 °C. Following biotinylation, free biotins were removed using the Ni-NTA column and biotinylated Ecads bound to Ni-NTA beads were eluted using a pH 7.5 buffer containing 200 mM Imidazole, 20 mM Na_2_HPO_4_, 500 mM NaCl, and 1 mM CaCl_2_.

### Single molecule AFM experiments

Purified canine Ecad monomers were immobilized on AFM cantilevers (Hydra 2R-50N; AppNano) and glass coverslips (CS) as described previously ^35,36^. Briefly, the CS and cantilevers were cleaned with 25% H_2_O_2_/75% H_2_SO_4_ overnight and washed with Milli-Q water. The CS was then cleaned with 1 M KOH and washed with Milli-Q water. Both the CS and cantilevers were washed with acetone and functionalized using 2% (v/v) 3-aminopropyltriethoxysilane (Millipore Sigma) solution dissolved in acetone. N-hydroxysuccinimide ester functionalized PEG spacers (MW 5000, Lysan Bio) were covalently attached to the silanized AFM tip and coverslip (100 mg/mL in 100 mM NaHCO_3_ dissolved in 600 mM K_2_SO_4_, for 4 h); 10% of the PEG spacers were decorated with biotin groups. Prior to a measurement, the functionalized AFM cantilever and coverslip were incubated overnight with BSA (1 mg/ml) to further reduce non-specific binding. The tip and surface were then incubated with streptavidin (0.1 mg/ml for 30 min; Thermo Fisher) and biotinylated canine Ecads (200 nM for 1h) were attached to the streptavidin. Finally, the surfaces were incubated with 0.02 mg/ml biotin for 10 min to block the free biotin binding sites on streptavidin.

Force measurements were performed using an Agilent 5500 AFM with a closed loop scanner. The force measurements were performed in a pH 7.5 buffer containing 10 nM Tris-HCl, 100mM NaCl, 10mM KCl and 2.5mM CaCl_2_. Cantilever spring constants were measured using the thermal fluctuation method ^37^. Unbinding events, which were characterized by stretching of the PEG tethers, were fit to a WLC model using a least square fitting protocol and specific events were chosen by discarding events that had a root mean squared error (RMSE) greater than the mean plus one standard deviation of all RMSEs. Additionally, since full-length PEG has a contour length of approximately 30 nm, events with contour lengths less than 30 nm were excluded. Furthermore, persistence lengths were constrained to 0.1 to 1 nm.

### MD Simulations and Analysis

MD simulations were performed with GROMACS 2020.1 using the FARM high-performance computing cluster at University of California, Davis as described previously ^35^. Briefly, the Ecad crystal structures in the absence of 19A11 (PDB ID code 2O72) and in the presence of 19A11 (PDB ID code 6CXY) were equilibrated by performing 60-ns molecular dynamics (MD) simulations. The protein structure was placed in the center of a dodecahedral box such that no atom of the protein was closer than 1 nm to any boundary. The box was solvated by adding water molecules and charge-neutralized by adding ions (150 nM NaCl, 4mM KCl, and 2mM CaCl_2_). Each simulation system contains ∼92500 atoms. The systems were relaxed using energy minimization, and stabilized by equilibrating under isothermal–isochoric and isothermal–isobaric conditions. Following equilibration, a 60-ns MD simulation was performed with 2-fs integration steps. The structures equilibrated after 20 ns, which was monitored by calculating the RMSD of the structures relative to the initial structure. The C-α RMSF of each residue in the Ecad EC1 domain (residue 1-100) during the final 10ns MD was calculated using *gmx rmsf* module. The distances between charged atoms for the salt bridge E13-R99 and K14-D58 were calculated using *gmx pairdist* module.

### SMD Simulations and Analysis

The last frame of each MD simulations was placed in the center of a rectangular box such that interacting Ecads were parallel with the longest axis of the box and no atom of the protein was closer than 1 nm to any boundary (30×12×8 nm for the 0ab conditions; 30×15×15 nm for the 1ab/2ab conditions). Each simulation system contained ∼380000 atoms for the 0ab conditions and ∼880000 atoms for the 1ab/2ab conditions. The simulation system was relaxed and equilibrated using the same methods as the MD simulations without isothermal-isochoric condition.

The change in Ecad interfacial area was estimated by the change in the solvent accessible surface area, which was calculated using *gmx sasa* module. The distances between center of mass of W2 and the corresponding pocket (residues 22-28, 36, 78-80, 89-92) were obtained using *gmx pairdist* module.

## Acknowledgments

Research in the SS lab was supported by the National Institute of General Medical Sciences of the National Institutes of Health (R01GM121885 and R01GM133880). Research in the BMG lab was supported by the National Institute of General Medical Sciences of the National Institutes of Health R35GM122467 (MIRA). The X-ray crystallography work was supported by National Institutes of Health/National Institute of Allergy and Infectious Diseases (contract no. HHSN272201700059C to PJM). This research used resources of the Advanced Photon Source, a U.S. Department of Energy (DOE) Office of Science User Facility operated for the DOE Office of Science by Argonne National Laboratory under Contract No. DE-AC02-06CH11357. Use of the LS-CAT Sector 21 was supported by the Michigan Economic Development Corporation and the Michigan Technology Tri-Corridor (Grant 085P1000817).

