## Supplementary Information for "Molecular mechanism for strengthening E-cadherin adhesion using a monoclonal antibody"

**Affiliations:** [1] Biophysics Graduate Group, University of California, Davis, CA; [2] Department of Biomedical Engineering, University of California, Davis, CA; [3] Seattle Children's Research Institute, Center for Developmental Biology and Regenerative Medicine, Seattle, WA; [4] Department of Biochemistry, University of Washington, Seattle, WA; [5] Seattle Structural Genomics Center for Infectious Disease (SSGCID), Seattle, WA; [6] UCB Pharma, Bainbridge Island, WA; [7] Center for Global Infectious Disease Research, Seattle Children's Research Institute, Seattle, WA; [8] Department of Pediatrics, University of Washington, Seattle, WA.

**Tables 1. Data collection and refinement statistics**

| | |
|---|---|
| Beamline | APS 21-ID-F |
| Space group | C2 |
| Cell dimensions | |
| a, b, c (Å) | 122.68, 77.47, 110.94 |
| α, β, γ (°) | 90.000, 92.905, 90.000 |
| Resolution (Å) | 50.0–2.20 (2.26–2.20) |
| No. reflections | 221,388 (16,425) |
| No. unique reflections | 52,838 (3,868) |
| Rmeas | 0.094 (0.672) |
| Rmerge | 0.082 (0.587) |
| I/σ(I) | 13.40 (2.60) |
| CC1/2 (%) | 99.7 (78.4) |
| Completeness (%) | 99.9 (99.9) |
| Redundancy | 4.2 (4.2) |
| Refinement | |
| Resolution (Å) | 40.09–2.20 |
| No. reflections | 52,833 |
| Rwork / Rfree | 0.1626 / 0.1970 (0.2370 / 0.3102) |
| No. atoms | |
| Protein | 4,849 |
| Ligand/ion | 90 |
| Water | 646 |
| B factors | |
| Protein | 35.04 |
| Ligand/ion | 56.07 |
| Water | 43.29 |
| R.m.s. deviations | |
| Bond lengths (Å) | 0.007 |
| Bond angles (°) | 0.892 |
| Ramachandran | |
| Preferred (%) | 97.91 |
| Allowed (%) | 1.93 |
| Outliers (%) | 0.16 |

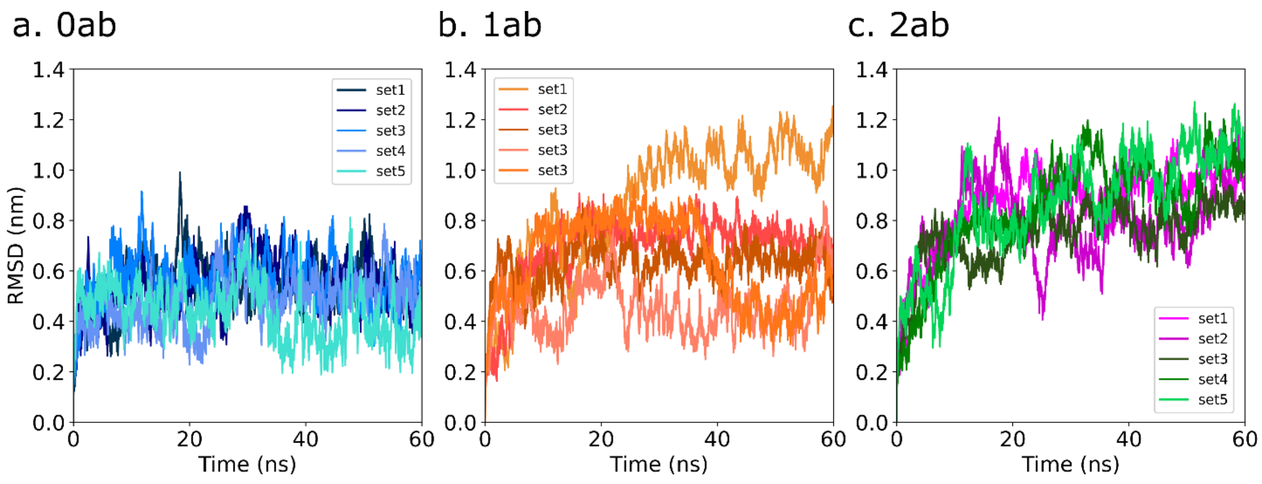


**Fig. 1**. RMSD of every simulation frame relative to the structures at the start of the simulations during the MD simulation. RMSD values observed in the (a) 0ab conditions, (b) 1ab conditions, and (c) 2ab conditions. RMSD values stabilized after 25ns for all simulations, suggesting the structures are well equilibrated.

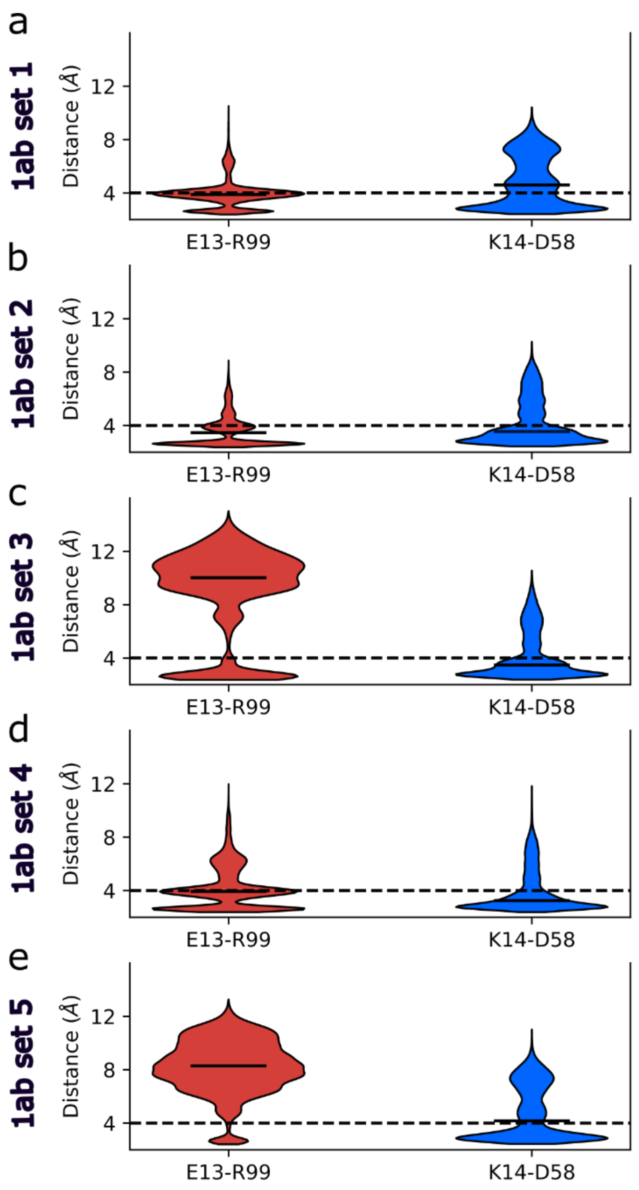


**Fig. 2. Formation of the E13-R99 and/or K14-D58 salt bridges during the 1ab conditions**. Violin plots for the distances between charged atoms measured in the E13-R99 and K14-D58 salt bridges observed in the last 40 ns of each MD simulation. (a) set 1, (b) set 2, (c) set 3. All of the simulation sets have at least one salt bridge formed during the MD.

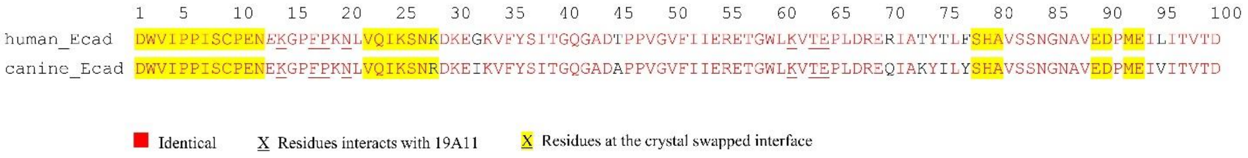


**Fig. 3**. Sequence comparison between human Ecad and canine Ecad reveals 91% identities.

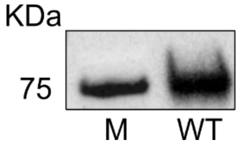


**Fig. 4**. Western-blot of Ecad with 19A11 primary antibody shows 19A11 binds to canine wild-type (WT) Ecad. Molecular ladder (M) with molecular weight 75KDa is shown.

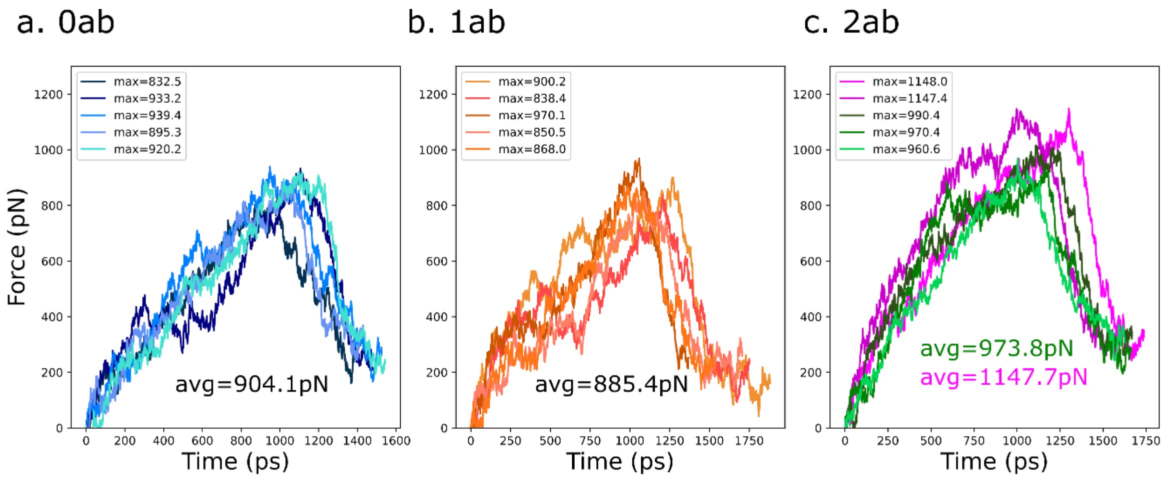


**Fig. 5**. Forces recorded during constant velocity SMD simulations in the (a) 0ab condition, (b) 1ab condition, and (c) 2ab condition. The average maximum forces observed in the 0ab and 1 ab condition are similar with values of 904.1 pN and 885.4 pN respectively. However, there are two populations observed in the 2ab conditions: set1/set2 is significantly higher with a value of 1147.7pN, while set3/set4/set5 is weaker with a value of 973.8pN.

**Supplemental Movie 1.** Example SMD simulations. One SMD example from each condition: 0ab (set 1), 1ab (set 4), 2ab weak conformation (set 5), and 2ab strong conformation (set 2) are shown. Ecad strand-swap dimers in the 0ab, 1ab and 2ab weak conformation break significantly faster compared with the 2ab strong conformation. Color scheme: Ecad (cyan and green), 19A11 heavy chain (magenta), and 19A11 light chain (orange).
